# Mechanical and kinetic factors drive sorting of F-actin crosslinkers on bundles

**DOI:** 10.1101/493841

**Authors:** Simon L. Freedman, Cristian Suarez, Jonathan D. Winkelman, David R. Kovar, Gregory A. Voth, Aaron R. Dinner, Glen M. Hocky

## Abstract

In cells, actin binding proteins (ABPs) sort to different regions in order to establish F-actin networks with diverse functions, including filopodia used for cell migration, or contractile rings required for cell division. Recent experimental work uncovered a passive mechanism that may facilitate spatial localization of ABPs: binding of a short crosslinker protein to two actin filaments promotes the binding of other short crosslinkers and inhibits the binding of longer crosslinkers (and vice versa). We hypothesize this sorting arises because F-actin is semiflexible and cannot bend over short distances. We develop a mathematical theory and a kinetic Monte Carlo simulation encompassing the most important physical parameters for this process, and use simulations of a coarse-grained but molecularly explicit model to characterize and test our predictions about the interplay of mechanical and kinetic parameters. Our theory and data predict an explicit dependence of crosslinker separation on bundle polymerization rate. We perform experiments that confirm a dependence on polymerization rate, but in an unanticipated non-monotonic manner. We use simulations to show that this non-monotonic behavior can arise in situations where crosslinkers have equal bundling affinity at equilibrium, but differing microscopic binding rates to filaments. This dependence of sorting on actin polymerization rate is a non-equilibrium effect, qualitatively similar to non-equilibrium domain formation in materials growth. Thus our results reveal an avenue by which cells can organize molecular material to drive biological processes, and can also guide the choice and design of crosslinkers for engineered protein-based materials.

---

Networks formed from filamentous actin polymers (F-actin) perform diverse mechanical tasks throughout cells, such as enabling migration (1, 2), adhesion (3), mechanosensing (4) and division (5). F-actin is formed into a network by crosslinkers, actin binding proteins (ABPs) that link multiple filaments. A large variety of crosslinkers exist, with diverse kinetic and mechanical properties (5). For example, the actin crosslinker fimbrin can bundle branched F-actin at the leading edge of migrating cells so that they can harness energy from actin polymerization to generate protrusive forces (1, 6). The force propagating F-actin cables that maintain a cell’s shape, or which are contained within a cytokinetic ring, each use their own F-actin crosslinking protein to form a specific geometry (7). Many ABPs may be involved in one single cellular mechanism. The cytokinetic ring of fission yeast employs formins to assemble F-actin, the crosslinker *α*-actinin to connect F-actin into anti-parallel bundles, and myosins to contract the bundles and ultimately divide the cell (8, 9). ABP kinetics can play subtle roles in these processes. For example, we previously showed that having optimal kinetics of binding (*k_on_*, *k_off_*), in addition to a an optimal binding affinity (*K_d_* = *k_off_*/*k_on_*) for the crosslinker *α*-actinin is crucial for proper contractile ring formation and constriction during cell division (10).

Regulating the spatial and temporal organization of ABPs in a crowded cellular environment is understandably complex, and determining the mechanisms involved is an active area of research. Some of this regulation may require explicit signaling pathways; for example generation of branched networks by the Arp2/3 complex can be activated by upstream activation of a Rho GTPase (11, 12). In addition to these signaling-based mechanisms, emerging data detail many *passive* mechanisms by which competition between different components for the same substrate can allow self-regulation and localization of ABPs in the actin cytoskeleton (13–16). We recently showed that *α*-actinin and fascin, two F-actin crosslinkers that are primarily found separated into different F-actin networks within cells, can self-sort in a simplified *in vitro* reconstitution of a branched Arp2/3 complex-nucleated network, and even sort to different domains within the same two-filament actin bundle (Figure 1A) (15). An outstanding challenge is to determine which of the biochemical characteristics of actin, fascin, and *α*-actinin yield sorting, and in that way determine if this mechanism may be generalizable to other polymers or crosslinkers.

**Fig. 1.**
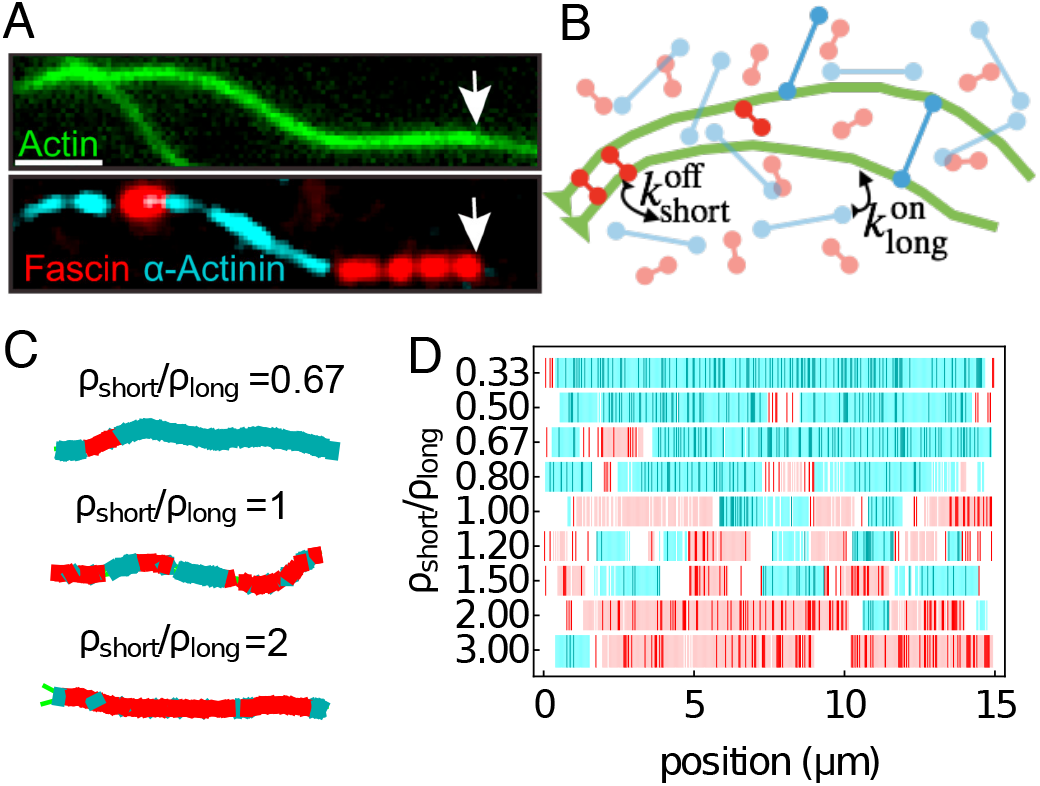
Crosslinker segregation in experiment and simulations. (A) Experimental three-color TIRF microscopy image showing two crosslinkers, fascin (red) and *α*-actinin (cyan) in domains on a two-filament actin bundle (green, arrows indicate polymerizing barbed end). Scale bar is 2 *μ*m. Adapted from Ref. (15). (B) Schematic of AFINES simulation: two filaments (green bead spring chains) are combined with two populations of crosslinkers, short (red) and long (cyan) that are represented as Hookean springs which can dynamically bind and unbind from filaments. Crosslinker opacity is proportional to number of heads bound (0,1, or 2). (C) Examples of steady-state filament bundles formed in AFINES, when two 15 μm filaments are mixed with an ensemble of long crosslinkers (*l*_long_ = 300 nm, shown in cyan) and short crosslinkers (*l*_short_ = 200 nm, shown in red), at different density ratios. The total crosslinker density is fixed at 0.5 *μ*m^-2^. See also Supp. Mov. 1. (D) Results from domain calculations for different density ratios. Cyan (red) lines show discretized position of long (short) crosslinkers while blue (pink) regions show extracted domains (see Materials and Methods and Figure S1). White regions are gaps between domains.

An important difference between fascin and *α*-actinin is their size; fascin is small (~8 nm), and therefore forms tight bundles composed of narrowly-spaced actin filaments, while *α*-actinin is larger (~35 nm) and therefore forms bundles composed of actin filaments that are more widely spaced (15, 17, 18). While filaments in *α*-actinin bundles are arranged with mixed polarity, fascin assembles bundles composed exclusively of parallel filaments, such that their fast-growing barbed ends all face the same direction (19, 20). Therefore the structures observed in our previous work (such as the one shown in Figure 1A) are parallel two-filament bundles in which the spacing between filaments alternates between approximately 8 and 35 nm (15). For transitions in bundle spacing, the actin filaments must bend significantly over length scales shorter than their persistence length *L_p_* = 17 *μ*m (21), which is energetically unfavorable. Since we observe domains in experiment, the energetic cost of bending must be compensated by favorable effects, such as the benefit of binding more crosslinkers and the entropic gain of mixing components on the bundle.

In this work, we use this hypothesis to develop a theoretical model that enables investigating the full range of mechanical and kinetic crosslinker properties that may lead to domain formation in F-actin bundles. We test this model in equilibrium systems with constant-length actin filaments using coarse-grained simulations and examine how the lengths of crosslinkers and the flexibility of F-actin affect crosslinker segregation. For non-equilibrium systems with growing filaments, our theoretical analysis predicts that actin polymerization and bundling affinity work together to determine the size of domains. We refine this prediction based on results from new *in vitro* experiments as well as coarse-grained simulations, and conclude that in addition to bundling affinity, the affinity of crosslinkers for single filaments changes how domains are formed. Thus, our theoretical models explain our experimental observations and elucidate passive mechanisms for crosslinker sorting in both equilibrium and non-equilibrium environments.

## Results and Discussion

### A. A simulation framework for actin and crosslinkers exhibits domain formation

Throughout this work we use AFINES (Active Filament Network Simulation), a coarse-grained molecular dynamics simulation framework built specifically for actin and ABP assemblies to investigate the mechanical properties of actin filaments and crosslinkers that might yield domain formation (schematic in Figure 1B and model details in SI: AFINES simulation) (22, 23). Actin filaments are modelled as polar worm-like chains (represented as beads connected by springs) that can grow from their barbed end by increasing the rest length of the barbed end spring at a constant rate, and adding a bead when that rest length is above a threshold (as done e.g. in Ref. (24)). Filaments repel each other via a soft (Hookean) excluded volume interaction. Crosslinkers are modelled as Hookean springs with two ends (heads) that can stochastically bind and unbind from actin filaments via a Monte-Carlo procedure that preserves detailed balance. To impose that crosslinkers enforce parallel (or anti-parallel) alignment of actin filaments, we include a harmonic bending energy term for crosslinkers, such that crosslinkers prefer to be bound perpendicularly to actin filament segments (see SI: AFINES simulation). The simulation proceeds in 2D via Brownian dynamics, without volume exclusion between the crosslinkers, to enable efficient sampling.

With this minimal mechanical-kinetic parameterization, our model exhibits segregation of different length crosslinkers on F-actin bundles, similar to those seen in experiment (Figure 1C). By discretizing the position of the crosslinkers doubly bound to actin filaments, and interpolating gaps between nearby crosslinkers, we can define domain boundaries in a manner consistent with experimental resolution (Figure 1D, details of domain length calculation in SI: Domain Length). Thus, we can use AFINES to systematically explore how specific crosslinker characteristics affect domain formation, and study domain formation in noisy conditions, in which bundling requires crosslinker-filament collision, and may be inhibited by actin filament fluctuations.

### B. Energetic cost of actin filament bending modulates domain formation

In our previous work, we modeled domain formation as a 1D process by which new crosslinkers are added to the barbed end of a growing two-filament bundle and do not unbind (15) (similar to models used for studying self-assembly of binary materials out of equilibrium (25, 26)). These assumptions were motivated by our experimental observations that bundling occurred at approximately the same rate as actin filament polymerization, and domain boundaries, once formed, remained fixed for the duration of the experiment (15). Using our model, we were able to fit an effective kinetic competition parameter, a kinetic barrier height of *ϵ* = 4.8*k_B_T*, to our experimental results (where *k_B_* is Boltzmann’s constant, and *T* is temperature) (15). Since we defined *ϵ* based on an Arrhenius-type equation, 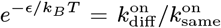, our predicted barrier height corresponds to a rate for adding a different crosslinker and starting a new barbed-end domain 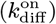 that is about 120-fold slower than the rate for continuing the same domain 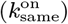; at equal concentrations, this resulted in long domains of both crosslinker types, as observed in experiments (15).

However, fitting this model does not make any connection to the underlying hypothesis that this kinetic competition is due to the cost of bending actin. To derive a similar model from first principles, we estimate the cost of bending actin such that, e.g., an *α*-actinin can be inserted with a gap length of *l_g_* along the filament from a fascin domain. As shown in the diagram in Figure 2A, the filament must bend twice at an angle *θ* for the filament bundle to switch domains (this geometry, where one filament is straight and the other bent, is based on cryoelectron microscopy images of domain switches in Ref. (15)). In the absence of filament fluctuations, we estimate *θ* ≈ arcsin (Δ*l_xl_*/*l_g_*) where Δ*l_xl_* is the difference in length between the two crosslinkers. Since the energy cost of bending an angle *θ* over a distance *l_g_* for a worm-like chain is *k_B_TL_p_θ*^2^/2*l_g_* (22, 27), where *L_p_* is the filament persistence length, *k_B_* is Boltzmann’s constant, and *T* is temperature, the total energy cost to bend a filament twice is

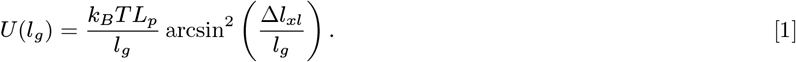

**Fig. 2.**
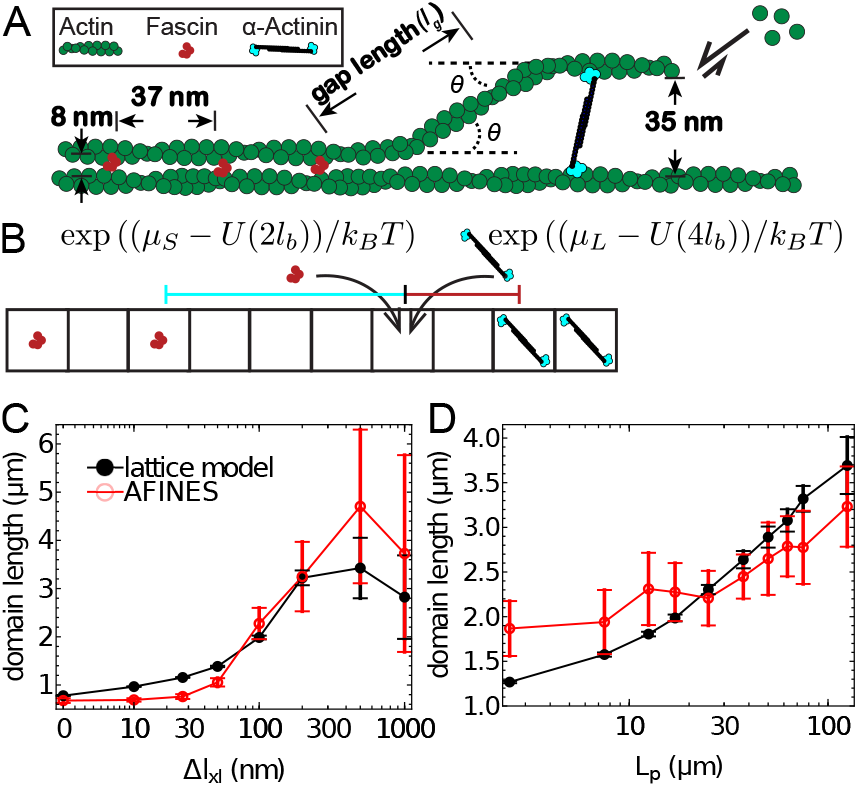
Effect of crosslinker and filament mechanics on domain length. (A) Schematic of a two filament bundle transitioning from a tight fascin bundle to a wider-spaced *α*-actinin bundle in an idealized geometry. The filament bends twice at an angle *θ* leaving a gap along the filament contour of length *l_g_*. (B) Schematic of the lattice model used to test energy term via Metropolis Monte-Carlo simulation. Exponential terms are the probabilities of the indicated lattice site switching from an unoccupied state to being occupied by a short (left) or long (right) crosslinker. *μ_L_* and *μ_S_* are the chemical potentials for the “long” and “short” crosslinker, and *l_b_* is the spacing between crosslinker binding sites in an actin bundle (see main text). (C) Domain lengths (averaged over 20 simulations; error bars are SEM) from AFINES and lattice model as a function of crosslinker length difference (Δ*l_xl_*), with *l*_short_ = 200 nm. (D) Same as C, but varying filament persistence length (*L_p_*). See also Supp. Mov. 2 and 3 for movies showing the effects in (C) and (D).

Equation (1) indicates that increasing the magnitude of mechanical parameters, such as the persistence length of filaments or the difference in length between crosslinkers, will increase the energy required to switch domain type on a filament bundle. The higher switching energy would decrease the likelihood of switching domains, and therefore increase domain lengths. While these mechanical characteristics are difficult to modulate experimentally, they are explicit parameters in AFINES, allowing us to directly test Equation (1) in simulation by exploring a range of crosslinker sizes and filament persistence lengths.

To quantitatively compare the results of our simulations with the predictions from Equation (1), we use an equilibrium 1D lattice model (Figure 2B) introducing the gap energy (Equation (1)) into a Monte Carlo (MC) simulation of a “bundle” of fixed length. The lattice contains a constant number of binding sites, *L*_0_/*l_b_* = 405, where *L*_0_ = 15 μm is the filament length and *l_b_* = 37 nm is the binding site spacing. Each lattice site can be in one of three states: empty, populated by a short crosslinker (S), or populated by a long crosslinker (L) (Figure 2B). The energy of the lattice is given by,

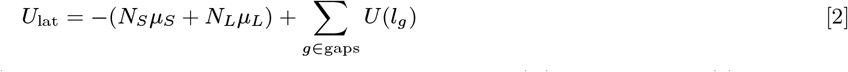

where *N*_*S*(*L*)_ is the number of short (long) crosslinkers, *μ*_*S*(*L*)_ are their chemical potentials, *U*(*l_g_*) uses Equation (1) and “gaps” is the set of all empty lattice patches between short and long crosslinkers.

We can measure *μ*_*S*(*L*)_ from our simulation data by determining the number of crosslinkers bound to two filaments in AFINES simulations with only one crosslinker (Figure S2). Assuming the energies in the corresponding lattice model are Boltzmann distributed, we can evaluate the partition function to determine the average lattice occupancy number 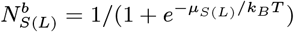, corresponding to 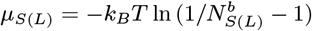. In the MC procedure, we compute the final state of the lattice using the Metropolis algorithm: iteratively, we switch a randomly chosen site to a randomly chosen new state with probability min (1, exp (−Δ*U*_lat_/*k_B_T*)), where Δ*U*_lat_ is the energy cost incurred by switching (28, 29).

To minimize the effects of kinetic factors in AFINES simulations, we fix the densities at *ρ*_short_ = *ρ*_long_ = 0.25 *μ*m^-2^, the actin filament length at *L*_0_ = 15 *μ*m, and the binding rates of the short and long crosslinkers, 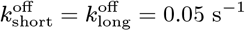 and 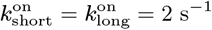. We find that for differences larger than 30 nm, domain length increases with Δ*l_xl_* (Figure 2C). Similarly, increasing the filament persistence length yields a significant increase in domain length (Figure 2D). We also observe that changes in filament stiffness can have a large effect on the rate of domain formation (see Supp. Mov. 3). Our results from the lattice simulation conform well with the AFINES simulations (Figure 2C-D) indicating that the mechanical model for crosslinker segregation is a good predictor for domain length in an equilibrium environment. This suggests that the driving forces controlling domain size are (1) the energetic cost of bending the actin, (2) the energy gain from binding, and (3) the entropy for mixing crosslinkers. Other factors, such as the entropy of filament fluctuations, and its dependence on gap size, do not appear to be major factors.

### C. Actin polymerization and crosslinker binding modulate domain formation in growing bundles

The mechanical parameters in Equation (1) are explicitly introduced into the simulations, and hence the dependence of domain size is easily testable. However, the average observed gap length at interfaces, 〈*l_g_*〉, is an emergent property within a bundle. In our experiments, the number of binding sites available to bind a new crosslinker can potentially be modulated by changing the polymerization rate of actin. Faster polymerization may promote generation of larger gaps before a crosslinker binds, and therefore a higher probability of switching. We demonstrate this behavior using a 1D lattice model, similar to that described above (Figure 2B) but with filament polymerization (schematic in Figure 3A). Polymerization is modeled by the steady addition of new sites for bundling at the barbed end (rightmost square in Figure 3A) of a 1D lattice. As sites become available, they can be filled by a crosslinker of the same type as the rightmost previous crosslinker with rate *k*_on_, and with the other type with rate *k*_on_*e*^-*U*(*l_g_*)/*k_B_T*^. Experiments show near simultaneous polymerization and bundling, with domain boundaries that do not change over time (15); we therefore include the approximation that crosslinkers remain fixed once bound, which enables extremely efficient calculations, while being a good proxy for what can be observed at experimental resolution. The dynamics of this lattice are simulated using a kinetic Monte Carlo (KMC) procedure, whereby the rates of all possible events (binding of either crosslinker to available sites, plus growth to add an additional available site) are computed, and then one event is selected randomly from this list weighted by the probability of that event occurring (29).

**Fig. 3.**
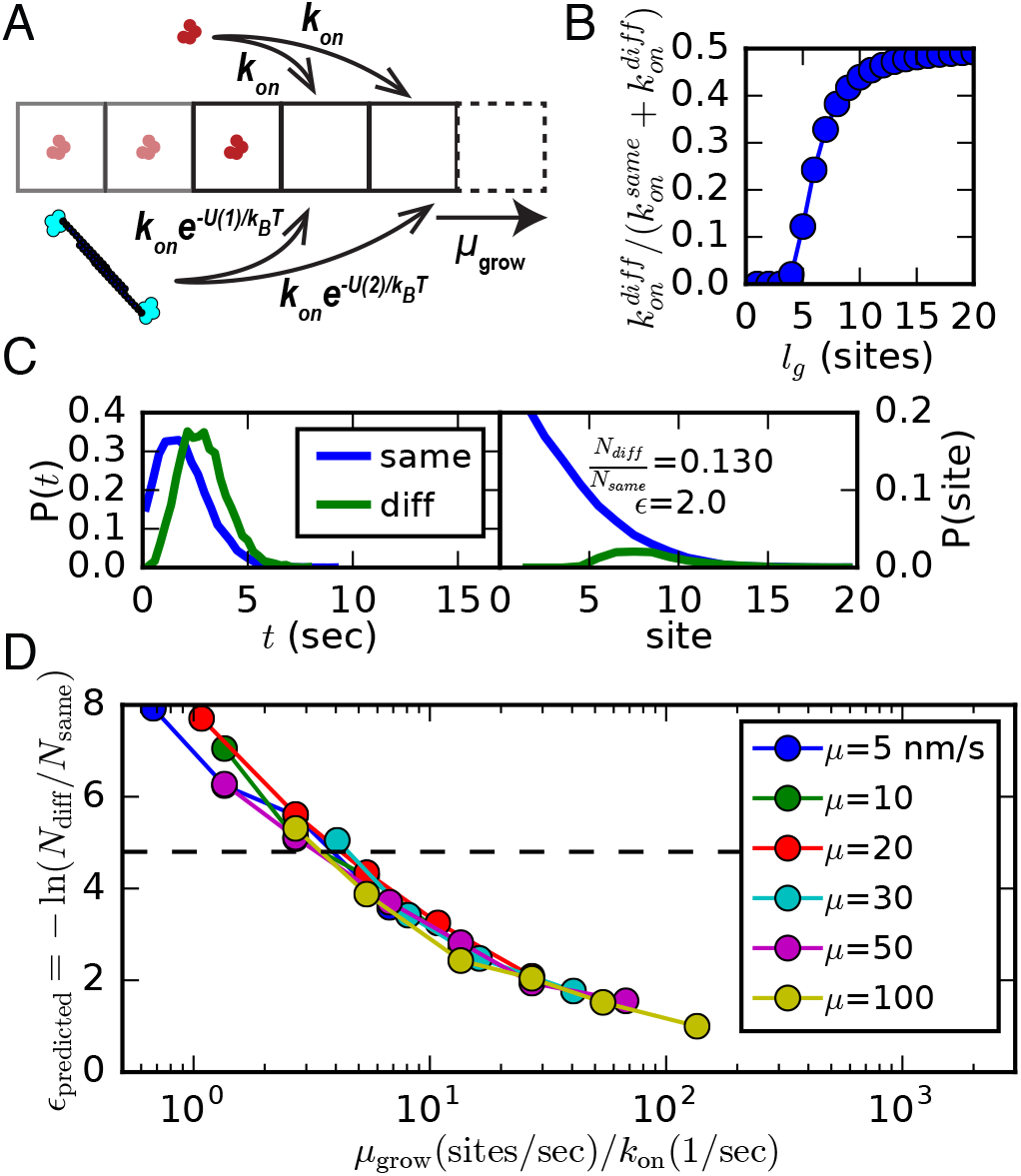
Energetic cost of filament bending leads to kinetic competition. (A) Numerical model for testing effect of filament growth on domain formation. New crosslinker binding sites are added at a rate *μ*_grow_, and crosslinkers are added via KMC at a rate governed by Equation (1). (B) Likelihood of adding the other (‘diff’) crosslinker at the barbed end at a given number of binding sites from the interface governed by Equation (1), assuming equal concentrations and affinity of crosslinkers. (C) Example results from the model in (A), with *μ*_grow_=100 sites/sec and *k*_on_=0.05/sec to a given site. Left: histogram of the amount of time that passes before binding the same or different crosslinker type. Right: histogram of how far from the current domain the next crosslinker adds. Inset: frequency of addition of switching bundle type, and the computed *ϵ* = − ln(*N*_diff_/*N*_same_). (D) Many experiments collapse onto one curve, showing that the amount of competition at the growing end depends only on the ratio of growth rate to bundling rate. The dashed line shows the value fit from experiments in Ref. (15). In (B-D), *L_p_* = 17*μ*m and Δ*l_xl_* = 27 nm. Each data point is an average over 10 simulations where the filament grows until it reaches a length of 15 *μ*m, and the kon values used range from 0.02/sec to 1.0/sec.

We test our model with *L_p_* = 17 *μ*m (corresponding to F-actin) and Δ*l_xl_* = 27 nm (corresponding to the difference in length between fascin and *α*-actinin). Figure 3B shows how the probability of a different crosslinker binding depends on the number of available sites. Interestingly, because the physiological crosslinker length difference is small, equal site binding probability is reached after only 20 binding sites (~ 0.7 *μ*m), much less than the persistence length of a filament (Figure 3B). As we would expect, if a switch of crosslinker type occurs, it generally requires a longer amount of time and occurs farther away from the existing domain than does a domain extension event (Figure 3C).

Given this, we hypothesized that it would be possible to modulate the amount of domain switching by changing the rate of bundling and the rate of filament polymerization. We scan realistic conditions for filament polymerization rate and binding rates and find that the amount of competition depends only on the ratio of these two quantities (Figure 3D). We find that the level of competition observed in Ref. (15) (dashed line in Figure 3D), corresponds to a polymerization rate that is four-fold faster than the binding rate to a single site; this result is consistent with the observation from TIRF experiments that the bundle is zipped up by crosslinkers at approximately the same rate as the actin polymerizes (15).

### D. Fascin and *α*-actinin domain lengths depend on actin polymerization rate

To test the prediction that polymerization rate affects the domain switching rate, and disentangle the role of kinetics in domain formation, we added monomeric actin to a mixture of fascin, and *α*-actinin, at two different actin concentrations, 0.75 *μ*M and 1.5 *μ*M (Figure 4A). The results (Figure 4B) showed that indeed, polymerization effects the size of observed domains. While we expected domain sizes of one or both to increase under slower actin polymerization conditions, we were surprised to see that *α*-actinin switches from the less-to more-dominant crosslinker. This suggests that kinetic effects can dominate over thermodynamic affinities, and make it advantageous to have one crosslinker on fast-growing bundles, and the other on slowly growing bundles.

**Fig. 4.**
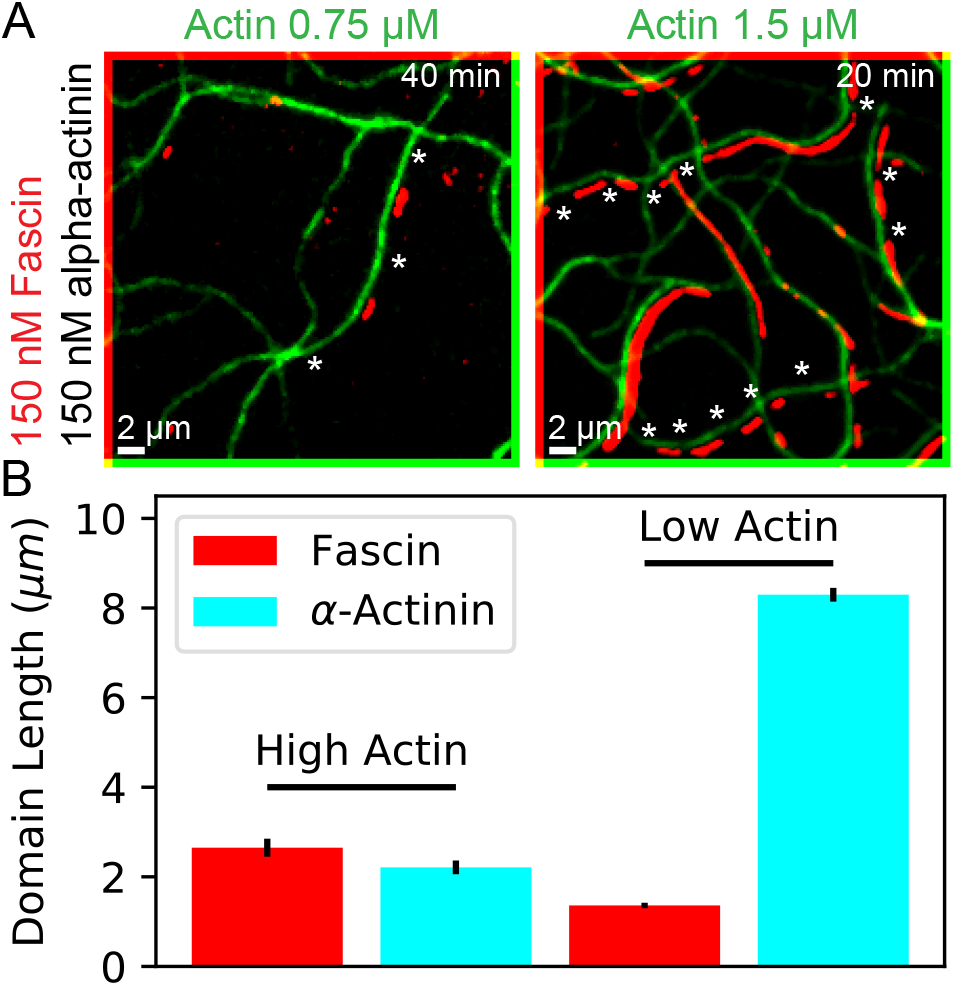
Kinetic competition in actin binding proteins. (A) Two-color TIRF image of actin filaments combined with fascin (labeled) and *α*-actinin (unlabeled) at two different actin concentrations (images diagonally offset by two pixels in each direction for clarity). *α*-actinin domains are inferred from regions with double actin fluorescence but no fascin (examples shown with stars). Supp. Mov. 4, and 5 correspond to these images. Experiments using labeled *α*-actinin and labeled fascin conform with these results but are harder to quantify (see Ref. (15), SI: TIRF data, and Figure S3). (B) Average domain length for fascin and *α*-actinin under these two actin concentrations. For each condition, two replicates were performed, and error bars are standard error of the mean domain size in each of the n=2 independent experiments.

Our previous work showed that both crosslinkers have similar bundling ability under these conditions, and fascin and *α*-actinin dissociated from actin bundles at nearly equal rates (15). Their affinity to a single actin filament is, however, different; we observe coating of single filaments by *α*-actinin, while we do not see significant residence of fascin on single filaments (see SI: TIRF data, movies). This may be due to the size and flexibility of *α*-actinin allowing it to bind to single filaments with both binding domains (18, 19). It is also possible that both crosslinkers have similar affinities to bundles, but the on- and off-rate constants themselves vary by a proportionality factor. The AFINES model allows us to directly test the hypothesis that kinetic factors can lead to a preference for one crosslinker domain on fast-growing filaments, and a preference for the other on slow-growing filaments.

### E. Simulation elucidates relationship between filament growth and binding kinetics

To investigate kinetic effects using AFINES, we first benchmark our simulations of growing filaments interacting with short and long ABPs against known experimental results (Figure 5A). Despite many simplifications in AFINES (including a lack of discrete binding sites and torsional restriction of actin) we obtain similar domain lengths as experiment with similar actin growth rates (40 nm/s), a short crosslinker length of *l*_short_ = 200 nm and long crosslinker length of *l*_long_ = 300 nm (Figure 5B). While it would be ideal to use crosslinkers that are the same length as in experiment, this was computationally infeasible in the simulation due to the required high spring stiffness (see SI: AFINES simulation for details); using shorter length differences yields the same characteristic highly cooperative domain length growth, but with shorter absolute domains (not shown).

**Fig. 5.**
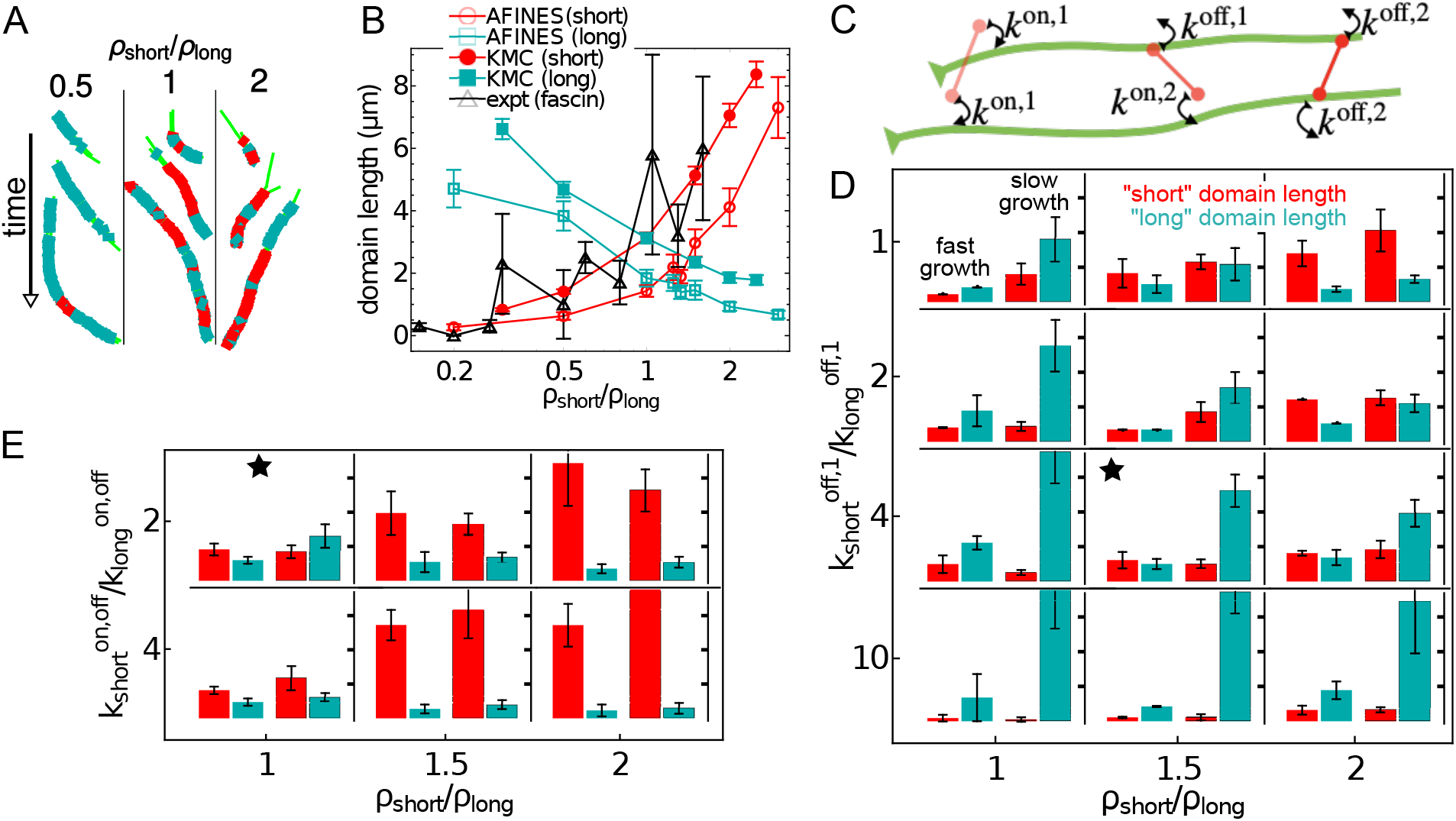
Competition between binding kinetics and polymerization in simulation. (A) AFINES trajectories of two filaments growing and forming crosslinker domains with a constant density of long (cyan) crosslinker (*ρ*_long_ = 0.25*μ*m^-2^) and varying the density of short (red) crosslinker (left to right: *ρ*_short_ = 0.125, 0.25, 0.5*μ*m^-2^, respectively). (B) Domain length as a function of density ratio for AFINES simulations of short and long crosslinkers, KMC simulations of two populations of crosslinkers, and experiments with actin, fascin, and *α*-actinin. For MC and AFINES simulations, *μ*_grow_ = 40nm/s with maximum filament length of 15*μ*m, and MC simulations used *k*_on_ = 0.26/s to a site, with each data point an average of 100 simulations. Experiments had comparable average polymerization rates for actin, data from Ref. (15). (C) Schematic of rate constant parameterization in AFINES (see SI: AFINES simulation for details). (D-E) Bar plots show domain lengths (ranging from 0 − 7 *μ*m, where each tick mark is 2 *μ*m) of short (red) and long (cyan) crosslinkers for fast actin polymerization (left, *μ*_grow_ = 100 nm/s) and slow actin polymerization (right, *μ*_grow_ = 10 nm/s) for varying kinetic conditions. Supp. Mov. 6 corresponds to (D,top-left) and best illustrates how kinetics effects domain size. Starred plots indicate conditions that most closely resembled experimental results (Figure 4B). (D) Varying the concentration *ρ*_short_ and single filament dissociation constant 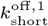 of the short crosslinker. (E) Varying the concentration *ρ*_short_ of the short crosslinker and rate constants 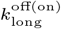 of the long crosslinker, such that 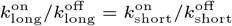. In (B,D,E), domain lengths are averaged over the 100 s after the filament reached its full length (*l*_max_ = 15 *μ*m) and 10 simulations, of which 5 were initialized with a short crosslinker bundle, and 5 were initialized with a long crosslinker bundle (SI: AFINES simulation); *ρ*_long_ = 0.25 *μ*m^-2^, *l*_short_ = 200 nm, *l*_long_ = 300 nm, and, unless otherwise specified, *k*^on^ = 2s^-1^ and *k*^off^ = 0.05s^-1^. Error bars are SEM.

In order to determine how crosslinker binding kinetics may effect domain formation on growing bundles, we relax our assumption that long and short crosslinkers have equal binding and unbinding rates to filaments. First, in order to assess our hypothesis that affinity for single filaments may differentiate *α*–actinin and fascin, we alter the equations determining binding for crosslinkers such that a crosslinker head may have one dissociation rate constant when bound to a single filament (*k*^off,1^) and another when both heads are bound (*k*^off,2^) as shown in Figure 5C (formula in SI: AFINES simulation). When crosslinkers have equivalent rate constants (Figure 5D, top row) faster growth yields shorter domains for both crosslinkers, consistent with our predictions from Figure 3. However, if the short crosslinker has lower single filament affinity (Figure 5D middle rows) then growth rate has the opposite effect; domain length remains constant or increases with growth rate, similar to the results observed in experiments for fascin (Figure 4). This behavior is magnified at higher densities of the short crosslinker. At very low single filament affinity (Figure 5D bottom row), domain formation is inhibited, indicating a threshold for the amount of time crosslinkers must remain bound to single filaments to form bundles.

We also use the simulation to test the effect of the short and long crosslinkers having different rate constants but equal (single filament and bundling) affinities, by keeping the ratio of their rate constants equal (i.e., 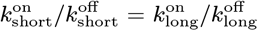). In this way we can test whether it is advantageous to be a fast-on/fast-off or a slow-on/slow-off crosslinker. We found that fast-on/fast-off crosslinkers typically yield larger domains than slow-on/slow-off (Figure 5E). Furthermore, at intermediate ratios of rates, fast filament growth may yield larger domains for the fast binding crosslinker than slow filament growth, as seen in experiment (Figure 5E, top row). However, very fast on/off rates yield larger domains at slower growth rates (Figure 5E, bottom row).

*These results indicate that the kinetics of crosslinker binding, and not only bundle affinity, are important for crosslinker segregation under non-equilibrium (polymerization) conditions*. Our observations from experiment that increasing growth rate decreased *α*-actinin (“long”) domain length but increased fascin (“short”) domain length correspond best with simulation when we decreased the single filament affinity of the short crosslinker slightly (Figure 5D, middle rows). A possible explanation for this result is that single filament affinity and crosslinker diffusion compete in bundle formation. In a slow growth environment, it is more advantageous to remain bound on a single filament than to freely diffuse, because the other (unbound) filament is likely nearby. In a fast growth environment, the unbound filament may bend away from the bound filament, so dissociating from the bound filaments enables the spatial exploration necessary to find an ideal bundling location. If affinities are equal, then higher rates of attachment and detachment are advantageous to forming larger domains (Figure 5E), perhaps because they allow for defects (crosslinker switches) at the growing front to anneal.

## Conclusions and Outlook

Mechanical properties of F-actin and ABPs are important for cytoskeletal function; for example, filopodia rely on the rigidity of F-actin bundles while actomyosin contractility depends on actin filament buckling (30, 31). Here we show a context in which these mechanical properties are also important for cytoskeletal self-organization. Specifically, our mechanical theory, and coarse-grained simulations highlight the importance of the length difference between crosslinkers and actin filament persistence length in forming segregated domains of crosslinkers on a single F-actin bundle. We predict that bundling proteins with larger length differences than *α*-actinin and fascin (27 nm) and polymers with larger persistence lengths than actin filaments (17 *μ*m) will have even more capacity to sort crosslinkers than previously observed. While it is experimentally difficult to control the length of a crosslinker independently of its binding affinity (32), engineered crosslinkers with tunable length spacers (for example, made from DNA) may enable controlled experiments in the future.

A possible limitation to our study is that the experimental setup may exhibit interactions between actin and the coverslip, which prevent crosslinkers from unbinding. Coarse-grained simulations do not have this limitation, and indeed exhibit domain flux, domain merging, and domain splitting, motivating future experiments on a passivated or lipid surface, where crosslinkers would be more free to unbind from the bundle.

Future modeling can expand on this work by incorporating further molecular details of actin filaments. For example, actin filaments have a helical structure, and binding at discrete sites requires aligning of the two helical pitches; this structural characteristic and the need for torsional strain on the actin may affect the binding length scale and spacing between crosslinkers of the same type. In this work, we have focused on the origin of sorting behavior in two-filament bundles previously observed in experiments. Simulations and experiments of cellular environments are necessary to determine if the specific mechanical and kinetic sorting principles studied here are sufficient to generate distinct actin network structures.

The cytoskeleton is highly dynamic, and there are many structures in the cell where the actin network is constantly turning over (1, 33). Furthermore all ABPs have characteristic times at which they bind, unbind, and perform their functions. These effects may be constrained by predictions based on non-equilibrium statistical mechanical principles primarily applied to self-assembly in materials (25, 26), and our simulation framework may provide new ways of testing those prior theories. Here, we have demonstrated that actin polymerization can strongly impact the amount of binding of crosslinkers in surprising and non-linear ways. In light of these results and the fact that cytoskeletal networks are constantly restructuring, we feel that the non-equilibrium effects on ABP binding and sorting merit further scrutiny.

## Materials and Methods

All data from this work is available from the authors upon request. Instructions and code for running and analyzing all simulations is available in the “xlink_sorting_paper” folder of https://github.com/Simfreed/AFINES (22, 23).

### TIRF Experiments

Actin was purified from rabbit muscle acetone powder and labeled on surface lysines with Alexa488-succinimidylester (Life Technologies, Grand Island, NY) as described in Refs. (34, 35). Human *α*-actinin-4, human fascin 1 were purified and labeleled with Cy5-Monomaleimide (GE Healthcare) or TMR-6-Maleimide (Life Technologies, Grand Island, NY) as described previously (15). Actin filament bundle lengths were measured using the ImageJ software (36). Additional experimental images can be found in SI: TIRF data, and experimental movies are also in the SI.

### Simulation domain length analysis

In AFINES, crosslinker can bind anywhere on filaments (i.e., they do not have discrete binding sites) and they do not have excluded volume interactions (i.e., their positions can overlap.) Therefore, to quantify domain length in simulation, we bin the crosslinkers by their position (relative to the top filament’s barbed end) into *l_b_* = 37 nm segments of the actin filament, corresponding to the size of an actin binding site (Figure S1). A segment is labeled “long” if it has more of the long crosslinker, “short” if it has more of the short crosslinker, and empty otherwise (Figure S1). For both the discretized AFINES simulation and the lattice models, we define a domain as a consecutive run of at least *n_d_* =4 crosslinkers separated by less than *l_d_* = 1 *μ*m, and its length is the distance between the first and last crosslinkers (Figure 1C and Figure S1). As in experiment, domain lengths were not calculated for the small set of simulations in which filaments did not form bundles (SI: AFINES simulation).

## ACKNOWLEDGMENTS

We thank members of the Dinner, Kovar, Voth, and Gardel labs for helpful conversations. This research was primarily supported by the University of Chicago Materials Research Science and Engineering Center (NSF Grant No. 1420709). Additional support was provided to G.M.H. by New York University. J.D.W. was supported by an NIH Ruth L. Kirschstein NRSA award (5F32GM122372-02). Simulation resources were provided by the University of Chicago Research Computing Center.

## Supporting Information

### AFINES simulation

In AFINES, actin filaments and passive crosslinkers are modeled as coarse-grained entities, whose positions are evolved in time using a combination of molecular dynamics and MC simulation. Actin filaments are treated as worm-like chains of *N*(*t*) + 1 beads, at time *t*, connected by *N*(*t*) harmonic springs (links) and *N* (*t*) − 1 angular harmonic springs. Thus, the internal forces on an actin filament can be obtained from the gradient of the potential energy *U_f_*(*t*):

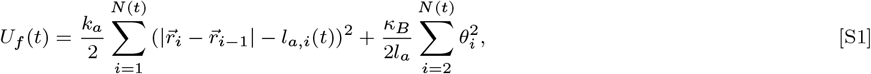

where 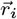 is the position of the *i^th^* bead on a filament, *θ_i_* is the angle between the *i^th^* and (*i* − 1)^*th*^ links, *k_a_* is the stretching force constant, *κ_B_* is the bending modulus, and *l_a,i_*(*t*) is the equilibrium length of a link. The persistence length of the filament is then *L_p_* = *κ_B_*/*k_B_T*, where *k_B_* is Boltzmann’s constant and *T* is the temperature.

The rest length of all the springs other than the one at the barbed end is constant, such that *l_a,j_*(*t*) = *l_a_* for *j* ∈ {2 … *N*(*t*)}. At every timestep the filament grows from the barbed end via an irreversible Markov process, such that

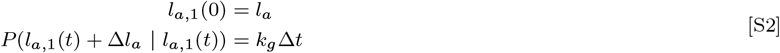

where *P*(*l_B_*|*l_A_*) is the probability of the barbed end link having length *l_B_* if on the previous timestep it had length A, Δ*l_a_* is the increase in segment length, *k_g_* is the rate of segment addition, and Δ*t* is the simulation timestep; *k_g_* Δ*l_a_* = *μ*_grow_ is thus the growth rate. If *l*_*a*,1_(*t*) + Δ*l_a_* > 2*l_a_*, the barbed end link is split into two links, such that each has rest length *l_a_*, and the total number of segments *N_a_*(*t*) is increased by 1. We set a maximum number of segments such that the filament stops growing when *N*(*t*) ≥ *N*_max_.

Springs on different filaments are restricted from overlapping via a soft (Hookean) excluded volume interaction,

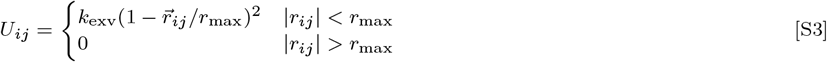

where *k*_exv_ is the interaction stiffness, *r_ij_* is the vector whose norm is the distance of closest approach between the springs *i* and *j*, and *r*_max_ is a cutoff distance for interaction. Excluded volume forces are propogated to filament beads via the lever rule defined below for crosslinkers (Equation (S5)).

We model crosslinkers as Hookean springs, with two ends (heads *L* and *R*) that can stochastically bind to and unbind from filaments. In addition to our previous implementation (22), we incorporate a bending term into the crosslinker, so that it prefers to bind perpendicularly to filaments, and have different binding rates for 1 and 2 heads. Thus, the potential energy of a crosslinker is

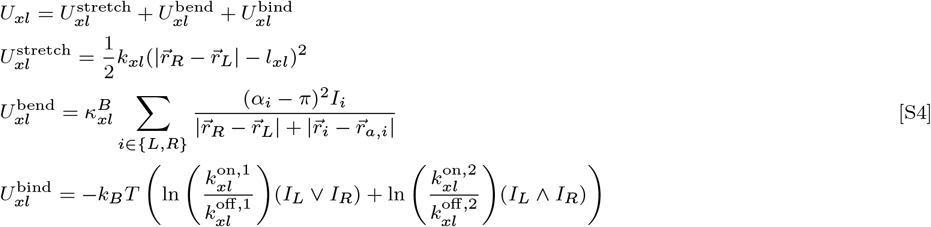

where *k_xl_* is the crosslinker stiffness, *l_xl_* is its rest length, 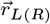 is the position of head *L*(*R*), *I*_*L*(*R*)_ is 1 if head *L*(*R*) is bound and 0 otherwise, 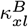 is the bending stiffness of the crosslinker, *α_i_* is the angle between the *i^th^* bound head and the filament spring to which it is bound, 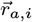 is the position of the nearest actin bead on that filament spring, 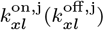 is the rate constant for a head binding (unbinding) if *j* − 1 heads (*j* ∈ {1, 2}) are already bound, and ∨ (∧) indicates the binary OR (AND) operation.

When a crosslinker is bound, it moves with the filament to which it is bound. When both crosslinker heads are bound, its tensile force, 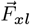 is propagated onto the filament beads neighboring each bound head at position 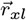 via the lever rule,

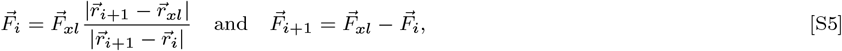

where 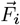 is the force on the filament bead at position 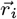.

Binding and unbinding are governed by an MC procedure constructed to satisfy detailed balance in the absence of motors. At each timestep of duration Δ*t*, an unbound crosslinker head becomes bound to the *i^th^* nearby filament with probability 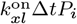, where *P_i_* is defined as follows. The closest point on the filament is identified, and the change in energy associated with moving the head to it, Δ*U_i_* is computed; *P_i_* = min [1, exp (−Δ*U_i_*/*k_B_T*)]. When a head becomes bound, its displacement, in the frame of reference of the filament link to which it attached, is stored as 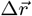. Later, the head can become unbound and displaced 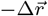 with probability 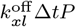, where *P* = min [1, exp(−Δ*U*/*k_B_T*)] and Δ*U* is the energy associated with the displacement. Because the dynamics depend on the allowed moves in the MC procedure, care must be used in interpreting the values of the rate constants. That said, the values that we obtain by tuning the parameters to yield behaviors consistent with experiments are generally within an order of magnitude of measured rate constants. As binding rate is affected by mechanical energy, and bending energy is added upon binding (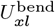 in Equation (S4)), our model exhibited a bias such that longer crosslinkers had higher affinity than shorter crosslinkers if they had the same 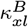; empirically, we found that setting 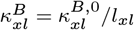 suppressed this bias (Figure S2).

We simulate the system using Brownian dynamics such that the position of an actin bead, motor head, or crosslinker head at time *t* is generated by the equation

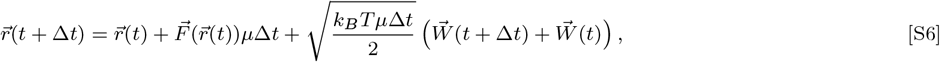

where 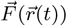 is the gradient of the potential of the particle, 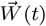 is a vector of random numbers drawn from the standard normal distribution, and we use the Stokes relation *μ* = 1/(6*πRν*) in the damping term, where *R* is the size of the particle, and *ν* is the dynamic viscosity of its environment (37). We simulate the system in 2D and use periodic boundary conditions to limit boundary effects. A complete list of model parameters used for Figures 1, 2 and 5 is provided in Table S1.

Guaranteeing that filaments formed bundles in simulation required very high crosslinker concentrations, which required significantly longer computation. In cases that filaments did bundle, we found that increasing crosslinker concentration above *ρ_xl_* = 0.25 *μ*m^-2^, did not significantly effect our domain length results. Thus, in our simulations we use *ρ_xl_* = 0.25 *μ*m^-2^, but promote forming a bundling via the initial conditions. Specifically, we initialize the two filaments to be parallel, at the same *x* coordinate, and separated in *y* by the average crosslinker length. Additionally we add a small bundle of crosslinkers (9 crosslinkers evenly spaced on the first filament link) at the barbed end of the filaments, to promote bundle formation. To prevent this initialization procedure from introducing bias toward one crosslinker, we run independent simulations of bundles initialized with a “short” crosslinker domain and a “long” crosslinker domain, and average over these simulations in our results (Figure 5C-E). A simulation was determined to have formed a bundle via the parallelness of the filaments; i.e., if for filaments 1 and 2, 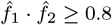, where 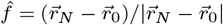 is the end to end vector of a filament, at all times in which domains are measured, then the filaments have bundled. As in experiment, simulations of actin filaments that did not yield bundles were not used to measure domain lengths.

### TIRF data

Additional results from TIRF microscopy can be found in Figure S3 and Supplemental Movies Figure 4, and Figure 5.

**Fig. S1.**
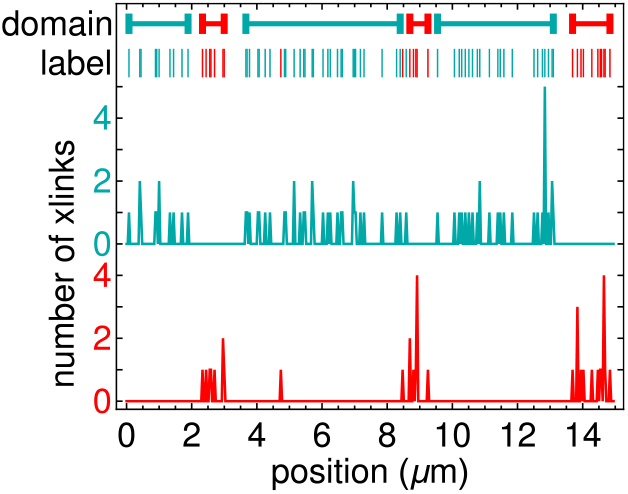
Domain calculation of a simulation with *ρ*_short_/*ρ*_long_ = 1, Δ*l_xl_* = 100 nm, and *L_p_* = 17 *μ*m at *t* = 1000 s. The distance (arc-length) of each crosslinker from the filament barbed end is calculated as the crosslinker’s position. Histograms of each crosslinker are calculated using a bin width of 37 nm, corresponding to the approximate size of F-actin binding sites. Each bin is then labeled with either a short crosslinker, long crosslinker, or nothing, depending on the majority. Domains are determined by iteratively merging nearby crosslinkers of the same type.

**Fig. S2.**
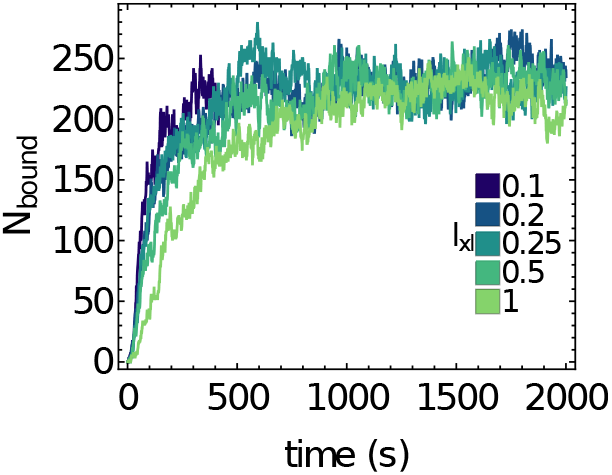
Number of crosslinkers doubly bound to a filament bundle for different crosslinker lengths have the same affinity, provided 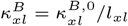. Each curve is a single simulation of two filaments bundling via one crosslinker.

**Table S1.** AFINES Parameter Values

| Sym-<br>bol | Description (units) [ref] | Value |
| --- | --- | --- |
| **Actin Filaments** |  |  |
| $N_f$ | number of filaments ($\mu\text{m}^{-2}$) | 2 |
| $N_{l,0}$ | initial number of links per filament | 2,15 |
| $N_{max}$ | max number of links per filament | 15 |
| $l_a$ | link rest length ($\mu\text{m}$) (38) | 1 |
| $k_a$ | stretching force constant ($\text{pN}/\mu\text{m}$) | 5 |
| $\kappa_B$ | bending modulus ($\text{pN}\mu\text{m}^2$) (21) | [0.01, 0.5] |
| $k_g$ | growth rate ($\text{s}^{-1}$) | 1 |
| $l_g$ | growth length ($\mu\text{m}$) | [0.008, 0.1] |
| $k_{\text{exv}}$ | excluded volume stiffness ($\text{pN}\mu\text{m}^2$) | 0.04 |
| $r_{\text{max}}$ | excluded volume cutoff ($\mu\text{m}$) | 0.05 |
| **Short Crosslinkers** |  |  |
| $\rho_{xl}$ | density ($\mu\text{m}^{-2}$) | [0.1, 0.75] |
| $l_{xl}$ | rest length ($\mu\text{m}$) (39) | 0.2 |
| $k_{xl}$ | stiffness ($\text{pN}/\mu\text{m}$) | 10 |
| $\kappa_{xl}^{B,0}$ | unnormalized bending stiffness ($\text{pN}\mu\text{m}^3$) | 0.04 |
| $k_{xl}^{\text{on},1}$ | maximum attachment rate from both unbound ($\text{s}^{-1}$) | 2 |
| $k_{xl}^{\text{off},1}$ | maximum detachment rate from one bound ($\text{s}^{-1}$) | 0.05, 0.2 |
| $k_{xl}^{\text{on},2}$ | maximum attachment rate from one bound ($\text{s}^{-1}$) | 2 |
| $k_{xl}^{\text{off},2}$ | maximum detachment rate from both bound ($\text{s}^{-1}$) | 0.05 |
| **Long Crosslinkers** |  |  |
| $\rho_{xl}$ | density ($\mu\text{m}^{-2}$) | [0.1, 0.4] |
| $l_{xl}$ | rest length ($\mu\text{m}$) (39) | [0.2, 1.2] |
| $k_{xl}$ | stiffness ($\text{pN}/\mu\text{m}$) | 10 |
| $\kappa_{xl}^{B,0}$ | unnormalized bending stiffness ($\text{pN}\mu\text{m}^3$) | 0.04 |
| $k_{xl}^{\text{on},1}$ | maximum attachment rate from both unbound ($\text{s}^{-1}$) | 0.5, 2 |
| $k_{xl}^{\text{off},1}$ | maximum detachment rate from one bound ($\text{s}^{-1}$) | 0.0125, 0.05 |
| $k_{xl}^{\text{on},2}$ | maximum attachment rate from one bound ($\text{s}^{-1}$) | 0.5, 2 |
| $k_{xl}^{\text{off},2}$ | maximum detachment rate from both bound ($\text{s}^{-1}$) | 0.0125, 0.05 |
| **Environment** |  |  |
| $\Delta t$ | dynamics timestep (s) | 0.00002 |
| $t_F$ | maximum simulated time (s) | 2000 |
| $X$ | length of assay ($\mu\text{m}$) | 35 |
| $Y$ | width of assay ($\mu\text{m}$) | 20 |
| $g$ | grid density ($\mu\text{m}^{-1}$) | 2.5 |
| $T$ | temperature ($K$) | 300 |
| $\nu$ | dynamic viscosity ($\text{Pa}\cdot\text{s}$) | 0.001 |
| $n_s$ | number of independent initializations (seeds) | 10, 20 |

**Fig. S3.**
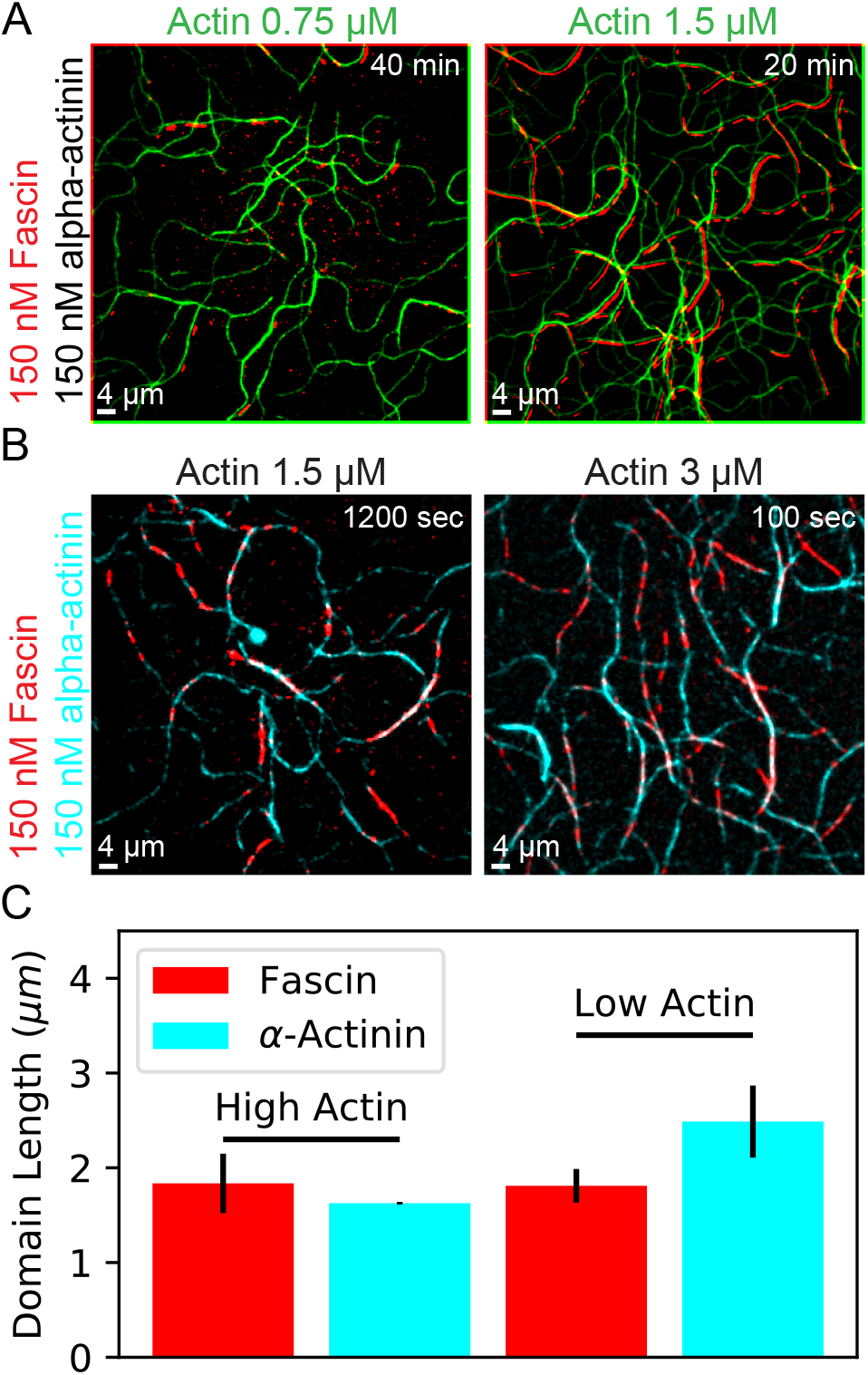
Additional TIRF data. (A) Wide field images of the same experiments shown in Figure 4. (B) Experiments using labeled *α*-actinin and labeled fascin, but unlabeled actin. Domains are evident here as well, however it is harder to quantify due to the binding of *α*-actinin to single filaments. Additionally it is harder to compare these results to Ref. (15), since the affinities of the crosslinkers to actin are different under these conditions. This experimental fact is evident comparing the amount of actin that polymerizes and bundles in (A,right), with (B,left). (C) Average domain length for fascin and *α*-actinin under these two actin concentrations, with actin unlabeled. The same qualitative trend is observed, where *α*-actinin becomes the dominant crosslinker under slower actin polymerization conditions. For each condition, two replicates were performed, and error bars are standard error of the mean domain size in each of the n=2 independent experiments.

**Supp. Mov. 1. AFINES Domain formation at different crosslinker density ratios**. AFINES simulations with (left-to-right) *ρ*_short_/*ρ*_long_ = {0.67, 1.0, 2.0}/*μ*m^2^, corresponding to the images in Fig. 1C. Hence, filament length is 15 *μ*m, *L_p_* = 17*μ*m, *l*_short_ = 200nm, and *l*_long_ = 300nm. *ρ*_short_ + *ρ*_long_ = 0.5.

**Supp. Mov. 2. AFINES Domain formation with differing crosslinker length differences**. AFINES simulations with (left-to-right) Δ*l*_xl_ = {10, 100, 500}nm corresponding to Fig. 2C. Hence, filament length is 15 *μ*m, *L_p_* = 17*μ*m, *l*_short_ = 200nm, and *ρ*_short_ = *ρ*_long_ = 0.25/*μ*m^2^.

**Supp. Mov. 3. AFINES Domain formation with differing filament persistence lengths**. AFINES simulations with (left-to-right) *L_p_* = {5, 25, 125}nm corresponding to Fig. 2D. Hence, filament length is 15 *μ*m, *l*_short_ = 200nm, *l*_long_ = 300nm, and *ρ*_short_ = *ρ*_long_ = 0.25/*μ*m^2^.

**Supp. Mov. 4. TIRF movie at low actin concentration**. TIRF movie corresponding to Fig. S3(A,left) with a zoom showing Fig. 4(A,left). Fascin is labeled in red, and actin is labeled in green. Color channels are offset by 2 pixels in both directions for clarity.

**Supp. Mov. 5. TIRF movie at high actin concentration**. TIRF movie corresponding to Fig. S3(B,left) with a zoom showing Fig. 4(B,left). Fascin is labeled in red, and actin is labeled in green. Color channels are offset by 2 pixels in both directions for clarity.

**Supp. Mov. 6. AFINES Domain formation with differing polymerization rates**. AFINES simulations with (left-to-right) *μ*_grow_ = {10, 100}nm/sec, corresponding to Fig. 5D, upper-left panel. Hence, filament length is 15 *μ*m, *L_p_* = 17*μ*m, *l*_short_ = 200nm, *l*_long_ = 300nm, and *ρ*_short_ = *ρ*_long_ = 0.25/*μ*m^2^.

## Notes

The authors declare no conflicts of interest.

## References

1. Pollard TD, Borisy GG (2003) Cellular motility driven by assembly and disassembly of actin filaments. Cell 112(4):453–465.

2. Lomakin AJ, et al. (2015) Competition of two distinct actin networks for actin defines a bistable switch for cell polarization. Nat. Cell Biol. 17(11):1435.

3. Parsons JT, Horwitz AR, Schwartz MA (2010) Cell adhesion: integrating cytoskeletal dynamics and cellular tension. Nat. Rev. Mol. Cell Biol. 11(9):633–643.

4. Engler AJ, Sen S, Sweeney HL, Discher DE (2006) Matrix elasticity directs stem cell lineage specification. Cell 126(4):677–689.

5. Lodish H, et al. (1995) Mol. Cell Biol. (WH Freeman New York) Vol. 3.

6. Welch MD, Iwamatsu A, Mitchison TJ (1997) Actin polymerization is induced by Arp 2/3 protein complex at the surface of listeria monocytogenes. Nature 385(6613):265.

7. Michelot A, Drubin DG (2011) Building distinct actin filament networks in a common cytoplasm. Curr. Biol. 21(14):R560–R569.

8. Vavylonis D, Wu JQ, Hao S, O’Shaughnessy B, Pollard TD (2008) Assembly mechanism of the contractile ring for cytokinesis by fission yeast. Science 319(5859):97–100.

9. Pollard TD (2010) Mechanics of cytokinesis in eukaryotes. Curr. Opin. Cell Biol. 22(1):50–56.

10. Li Y, et al. (2016) The F-actin bundler *α*-actinin Ain1 is tailored for ring assembly and constriction during cytokinesis in fission yeast. Mol. Biol. Cell. 27(11):1821–1833.

11. Rohatgi R, et al. (1999) The interaction between N-WASP and the Arp2/3 complex links Cdc42-dependent signals to actin assembly. Cell 97(2):221–231.

12. Goley ED, Welch MD (2006) The ARP2/3 complex: an actin nucleator comes of age. Nat. Rev. Mol. Cell Biol. 7(10):713.

13. Burke TA, et al. (2014) Homeostatic actin cytoskeleton networks are regulated by assembly factor competition for monomers. Curr. Biol. 24(5):579–585.

14. Suarez C, Kovar DR (2016) Internetwork competition for monomers governs actin cytoskeleton organization. Nat. Rev. Mol. Cell Biol. 17(12):799.

15. Winkelman JD, et al. (2016) Fascin-and *α*-actinin-bundled networks contain intrinsic structural features that drive protein sorting. Curr. Biol. 26(20):2697–2706.

16. Christensen JR, et al. (2017) Competition between Tropomyosin, Fimbrin, and ADF/Cofilin drives their sorting to distinct actin filament networks. Elife 6:e23152.

17. Jansen S, et al. (2011) Mechanism of actin filament bundling by fascin. J. Biol. Chem. pp. jbc–M111.

18. Hampton CM, Taylor DW, Taylor KA (2007) Novel structures for *α*-actinin: F-actin interactions and their implications for actin–membrane attachment and tension sensing in the cytoskeleton. J. Mol. Biol. 368(1):92–104.

19. Courson DS, Rock RS (2010) Actin crosslink assembly and disassembly mechanics for alpha-actinin and fascin. J. Biol. Chem. pp. jbc–M110.

20. Blanchoin L, Boujemaa-Paterski R, Sykes C, Plastino J (2014) Actin dynamics, architecture, and mechanics in cell motility. Physiol. Rev. 94(1):235–263.

21. Ott A, Magnasco M, Simon A, Libchaber A (1993) Measurement of the persistence length of polymerized actin using fluorescence microscopy. Phys. Rev. E 48(3):R1642–R1645.

22. Freedman SL, Banerjee S, Hocky GM, Dinner AR (2017) A versatile framework for simulating the dynamic mechanical structure of cytoskeletal networks. Biophys. J 113(2):448–460.

23. Freedman SL, Hocky GM, Banerjee S, Dinner AR (2018) Nonequilibrium phase diagrams for actomyosin networks. Soft Mat., In Press.

24. Zimmermann D, et al. (2017) Mechanoregulated inhibition of formin facilitates contractile actomyosin ring assembly. Nature Comm. 8(1):703.

25. Whitelam S, Hedges LO, Schmit JD (2014) Self-assembly at a nonequilibrium critical point. Phys. Rev. Lett. 112(15):155504.

26. Nguyen M, Vaikuntanathan S (2016) Design principles for nonequilibrium self-assembly. Proc. Natl. Acad. Sci. 113(50):14231–14236.

27. Rubinstein M, Colby RH (2003) Polymer Physics. (Oxford University Press, Oxford, United Kingdom).

28. Metropolis N, Rosenbluth AW, Rosenbluth MN, Teller AH, Teller E (1953) Equation of state calculations by fast computing machines. J. Chem. Phys. 21(6):1087–1092.

29. Newman M, Barkema G (1999) Monte Carlo Methods in Statistical Physics. (Oxford University Press).

30. Lenz M, Thoresen T, Gardel ML, Dinner AR (2012) Contractile units in disordered actomyosin bundles arise from f-actin buckling. Phys. Rev. Lett. 108(23):238107.

31. Murrell MP, Gardel ML (2012) F-actin buckling coordinates contractility and severing in a biomimetic actomyosin cortex. Proc. Natl. Acad. Sci. USA 109(51):20820–20825.

32. Li Y (2014) Ph.D. thesis (The University of Chicago).

33. Plastino J, Blanchoin L (2019) Dynamic stability of the actin ecosystem. J. Cell Sci. 132(4):jcs219832.

34. Isambert H, et al. (1995) Flexibility of actin filaments derived from thermal fluctuations. effect of bound nucleotide, phalloidin, and muscle regulatory proteins. J. Biol. Chem. 270(19):11437–11444.

35. Spudich JA, Watt S (1971) The regulation of rabbit skeletal muscle contraction i. biochemical studies of the interaction of the tropomyosin-troponin complex with actin and the proteolytic fragments of myosin. J. Biol. Chem. 246(15):4866–4871.

36. Schneider CA, Rasband WS, Eliceiri KW (2012) Nih image to imagej: 25 years of image analysis. Nat. methods 9(7):671.

37. Leimkuhler B, Matthews C (2013) Robust and efficient configurational molecular sampling via langevin dynamics. J. Chem. Phys. 138(17):174102.

38. Odijk T (1983) The statistics and dynamics of confined or entangled stiff polymers. Macromolecules 16(8):1340–1344.

39. Ferrer JM, et al. (2008) Measuring molecular rupture forces between single actin filaments and actin-binding proteins. Proc. Natl. Acad. Sci. USA 105(27):9221–9226.

